# Inferring unobserved vector dynamics for dengue forecasting using physics-informed neural networks and mechanistic transmission models

**DOI:** 10.64898/2026.02.13.705693

**Authors:** Jiayue Cao, Ziqiang Cheng, Min Su, Cang Hui

## Abstract

Accurately characterising mosquito infection dynamics is essential for effective dengue prevention and control, yet these dynamics are rarely observable through routine surveillance. Here, we integrate a reduced SI-SIR transmission model with monthly dengue incidence data using physics-informed neural networks (PINNs) to develop a Dengue-Informed Neural Network (DINN). The DINN simultaneously fits reported dengue cases from 15 countries between January 2014 to April 2025 and reconstructs the unobserved time series of infected mosquitoes. Using the inferred mosquito infection dynamics together with five-month-lagged climatic variables, we subsequently train recurrent neural networks (RNN, GRU and LSTM) to forecast mosquito infection levels one month ahead. For each country, the optimal forecasting model is selected based on out-of-sample performance, and model behaviour is interpreted using SHAP analysis. We show that the DINN captures multi-wave outbreak dynamics across diverse epidemic regions and enables the development of country-specific vector forecasting models. When integrated with the transmission model, the predicted mosquito trajectories support two-year projections of dengue incidence, providing a quantitative framework for early warning and evidence-based control strategies.

**Author summary:** Mechanistic models of dengue transmission often require assumptions about mosquito population sizes and temperature-dependent biological traits that are difficult to specify accurately and rarely observed in practice. These limitations can reduce the reliability of outbreak prediction and hinder the evaluation of control strategies. Here, we introduce a Dengue-Informed Neural Network (DINN) that integrates deep learning with a reduced SI-SIR transmission model to address these challenges. Without requiring prior specification of mosquito initial conditions or biological trait functions, DINN infers latent time series of infectious mosquitoes directly from reported human case data and reconstructs complex, multi-wave transmission dynamics. Applying this approach to dengue surveillance data from 15 countries, we uncover substantial regional variation in inferred mosquito infection patterns, reflecting differences in environmental variability and vector control. By combining the inferred mosquito dynamics with climatic information, we further develop interpretable forecasting models and generate two-year projections of dengue incidence. Our results demonstrate that DINN provides a flexible and interpretable framework for inferring unobserved vector dynamics and offers practical support for early warning and evidence-based control of mosquito-borne diseases.

## Introduction

The global incidence of dengue has increased substantially over recent decades, driven by factors such as climate change, urbanisation, and increased human mobility. According to the World Health Organization [1], reported dengue cases rose from 505,430 in 2000 to 14.6 million in 2024. As a vector-borne disease, dengue transmission is tightly coupled to the ecology and behaviour of mosquito vectors [2,3]. The primary global vector is *Aedes aegypti*, while *Aedes albopictus*, a secondary vector with greater tolerance to temperate climates, has contributed to outbreaks across an expanding geographic range [4]. In the absence of effective antivirals or widely accessible vaccines, dengue continues to impose substantial health and economic burdens, frequently overwhelming local healthcare systems during seasonal epidemics [5]. These challenges highlight the need for reliable early warning systems capable of supporting proactive vector control, efficient resource allocation, and timely public health interventions [6].

Mechanistic models, particularly parametric systems of differential equations, have long played a central role in characterising dengue transmission dynamics and informing control strategies [7-12]. For example, Huber et al. [13] quantified unimodal temperature responses of *Ae. aegypti* life-history traits using laboratory experiments and incorporated these relationships into transmission models to explore outbreak size, duration, and peak intensity under different climatic regimes. Building on this approach, Caldwell et al. [14] developed a climate-driven SEI–SEIR model that integrates the effects of temperature, humidity, and rainfall on mosquito physiology, successfully reproducing the timing and magnitude of dengue, chikungunya, and Zika outbreaks across distinct ecological settings in Ecuador and Kenya. Other studies have identified large-scale climate drivers, such as the Indian Ocean Basin-wide index, as potential predictors of dengue dynamics [15], or have incorporated periodic transmission parameters to capture local outbreak patterns [16]. Despite their interpretability and strong mechanistic grounding, these models often rely on fixed or laboratory-derived assumptions for key parameters (e.g., transmission rates or temperature response functions) that may inadequately capture the nonlinear and time-varying processes observed in real-world systems. As a result, model adaptability and predictive performance can be reduced, particularly across heterogeneous regions and during multi-wave seasonal outbreaks.

In parallel, data-driven methods based on machine learning have emerged as powerful tools for dengue forecasting and the development of early warning systems [18-23]. These methods have been widely applied to epidemiological time series prediction, climate-disease association analysis, spatiotemporal risk mapping, and public health decision support. Models such as Long Short-Term Memory (LSTM) networks, Support Vector Machines (SVM), Random Forests (RF), Gradient Boosting Machines (GBM), and hybrid architectures have demonstrated strong predictive performance by integrating meteorological, climatic, epidemiological, and geospatial data. For instance, Kakarla et al. [24] used an LSTM architecture incorporating relative humidity, soil moisture, temperature, precipitation, and the Niño3.4 index to predict dengue incidence in Kerala, India. Cheng et al. [25] proposed a hybrid framework combining Distributed Lag Nonlinear Models (DLNMs) with optimisation algorithms and machine-learning classifiers to analyse dengue outbreaks in southern China, while Ningrum et al. [26] applied ensemble learning approaches to predict outbreak occurrence and case counts in Indonesia. Lowe et al. [27] further demonstrated that incorporating seasonal climate forecasts and real-time ENSO indices into a Bayesian hierarchical framework improved predictions of outbreak timing in Ecuador. Despite their strong predictive performance, most data-driven models function largely as “black boxes”, offering limited interpretability and making it difficult to elucidate ecological mechanisms, quantify indirect environmental effects, or project transmission dynamics.

To bridge the gap between mechanistic interpretability and data-driven flexibility, recent studies have integrated epidemiological models into Physics-Informed Neural Networks (PINNs) [28-35]. PINNs embed governing equations directly into the neural network loss function, enabling simultaneous data fitting and equation-constrained learning. For example, Kharazmi et al. [28] applied PINNs to infer time-varying parameters and unobserved states in integer-order, fractional-order, and delay differential equation models of COVID-19. Millevoi et al. [33] introduced a split-training PINN framework that improved parameter estimation when incorporating hospitalization data into susceptible-infected-recovered (SIR) models. He et al. [32] proposed a Transmission Dynamics-Informed Neural Network (TDINN) that embeds a susceptible-exposed-infectious-recovered-dead (SEIRD) model within a neural network architecture to infer time-varying transmission rates across multiple pandemic waves. Their analysis revealed substantial lags between infection peaks and inferred transmission-rate dynamics, underscoring the importance of behavioural feedback and delayed intervention effects. Li et al. [35] extended this framework to multi-strain COVID-19 transmission, enabling joint inference of strain-specific parameters and competitive interactions among variants. Together, these studies demonstrate that PINNs can retain mechanistic interpretability while flexibly capturing time-varying dynamics under sparse data and complex, multi-wave epidemic patterns.

Despite these advances, PINN-based approaches have been applied predominantly to directly transmitted diseases, and their application to vector-borne diseases such as dengue remains limited. A central challenge is that key vector-related variables, most notably the number of infected vectors (e.g., mosquitoes), are typically unobserved in routine surveillance data. In this study, we address this limitation by developing a Dengue-Informed Neural Network (DINN) that integrates deep learning with a reduced SI-SIR transmission model. Building on PINN principles, DINN treats the temporal dynamics of infected mosquitoes as a latent neural network output, while embedding the residuals of the epidemiological equations into the loss function to constrain learning. This framework enables simultaneous fitting of reported human dengue cases and inference of unobserved infectious vector dynamics, without requiring prior specification of mosquito initial conditions or temperature-dependent biological trait functions.

We apply DINN to monthly dengue surveillance data from 15 countries to reconstruct multi-wave outbreak trajectories and infer region-specific dynamics of infectious mosquitos. Using the DINN-inferred mosquito time series, together with five-month-lagged temperature and precipitation data, we evaluate three recurrent neural network-based forecasting models: a vanilla Elman-type RNN, a gated recurrent unit (GRU), and a long short-term memory (LSTM) network, to predict mosquito infection level one month ahead. For each country, the optimal forecasting model is selected based on out-of-sample mean squared error. Forecasting model interpretability is further enhanced using Shapley Additive Explanations (SHAP) analysis [43]. Finally, the predicted mosquito dynamics are integrated into the DINN-parameterised reduced SI-SIR models to generate twenty-four-month forecasts of dengue incidence. Together, these results demonstrate that DINN provides a flexible and interpretable framework for inferring unobserved vector dynamics and supporting region-specific early warning and strategic planning for dengue control.

## Materials and methods

### Data Sources

Monthly reported dengue cases from January 2014 to April 2025 (*n*_*d*_ = 136 months) were obtained for 15 countries from the World Health Organization (WHO) [1]. The dataset, updated annually, provides the number of new cases reported in month *t*_*i*_ (*I*_new_(*t*_*i*_), where *i* = 1,…,*n*_*d*_), from which the cumulative number of reported cases up to month *t*_*i*_ was calculated as *I*_cum_(*t*_*i*_) = *I*_cum_(*t*_*i*_ ― 1) + *I*_new_(*t*_*i*_), with *I*_cum_(*t*_0_) = 0 at the start of the study period. To incorporate environmental drivers, we collected monthly mean temperature (*T*(*t*_*i*_)) and monthly total precipitation (*P*(*t*_*i*_)) for the same countries and time period from the National Oceanic and Atmospheric Administration (NOAA; S1 Fig) [36]. Monthly mean temperature for each country was computed as the average of daily temperature observations from all available meteorological stations within the country during that month, while the monthly total precipitation was obtained by summing the monthly precipitation totals reported by these stations. Annual population size data for each country were obtained from the World Bank and Worldometer [37,38].

### Reduced SI-SIR Transmission Model

Classical SI-SIR models are widely used for theoretical analyses of vector-born disease transmission [13-17]. In this framework, the human population at time t is divided into three compartments: susceptible *S*_*h*_(*t*), infected *I*_*h*_(*t*), and recovered *R*_*h*_(*t*). The total human population of a country at time t is denoted by

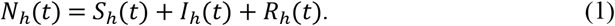

In addition, we introduce an auxiliary compartment, *I*_cum_(*t*), to track the cumulative number of confirmed dengue infections [35]. The mosquito population is similarly structured into susceptible *S*_*v*_(*t*) and infected *I*_*v*_(*t*) vectors.

Dengue transmission occurs from infected mosquitoes to humans, with new human infections arising through contacts between susceptible humans and infected mosquitoes. Environmental factors such as temperature and precipitation indirectly modulate transmission dynamics [2,9,13-15]. Specifically, mosquito life-history traits (e.g., lifespan and biting rate) are strongly temperature-dependent [2,13-15], while mosquito carrying capacity and mortality are sensitive to precipitation variability [14]. Accordingly, we define the vector-to-human transmission rate as,

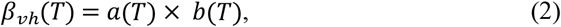

where *a*(*T*) is the temperature-dependent biting rate and *b*(*T*) is the probability of successful transmission following a bite by an infected mosquito. Both functions are modelled using Brière formulations [9,13-15]:

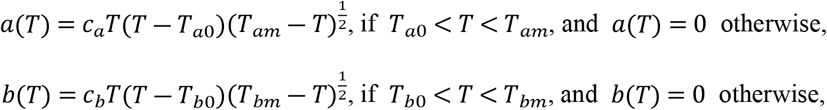

where *c*_*a*_, *c*_*b*_ are the rate constants, *T*_*a*0_ and *T*_*b*0_ are the critical thermal minima, and *T*_*am*_, *T*_*bm*_ are the critical thermal maxima, with their values taken from previous studies [13,14]: *c*_*a*_ = 2.02e-04, *c*_*b*_ = 8.49e-04, *T*_*a*0_ = 13.35, *T*_*b*0_ = 17.05, *T*_*am*_ = 40.08, *T*_*bm*_= 35.83.

In practice, however, mosquito infection data are rarely available and are often highly uncertain in real-world surveillance systems. To address this limitation, we employ a reduced SI-SIR framework [13,17], in which the number of infectious mosquitoes *I*_*v*_(*t*) is treated as a time-varying latent variable and inferred directly from human epidemiological observations (Fig 1).

**Fig 1.**
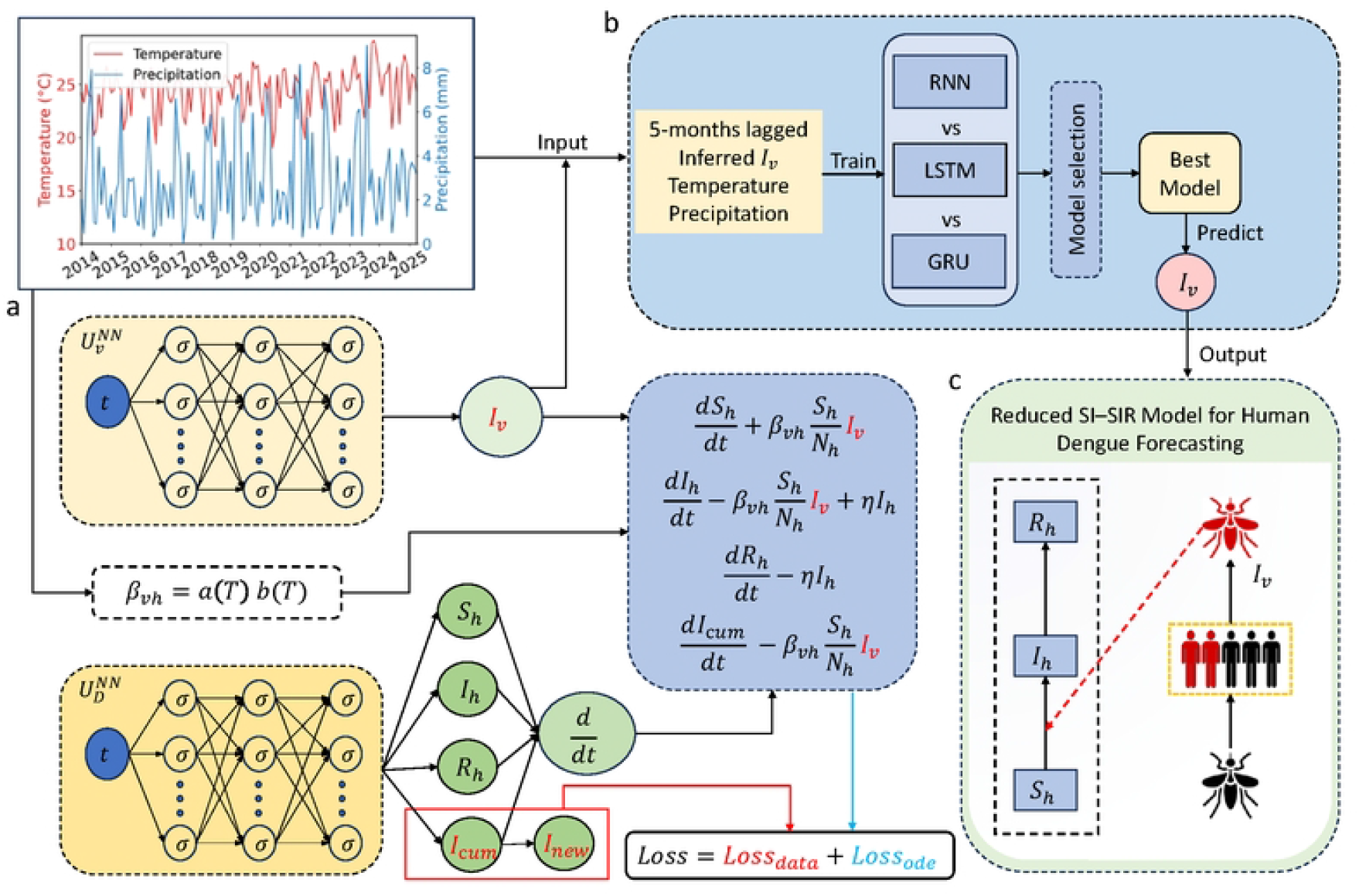
Schematic diagram of the DINN modelling and prediction framework. (a) Reconstruction of mosquito infection dynamics *I*_*v*_(*t*) from dengue surveillance data using DINN; (b) Selection of the optimal recurrent neural network (RNN, GRU, or LSTM) using five-month lagged mosquito infection and climatic variables; (c) Forecasting of human dengue incidence by integrating twenty-four-month predicted mosquito infection trajectories into the DINN-parameterised reduced SI-SIR model. Within DINN, 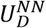 approximates human compartment states, and 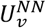 infers the latent infected-mosquito time series *I*_*v*_(*t*). The symbols σ and *d*/*dt* denote the activation function and automatic differentiation operator, respectively.

This reduced SI-SIR formulation captures dengue transmission between mosquitoes and humans while explicitly accounting for the limited observability of vector infection dynamics. The transmission model is governed by the following system of ordinary differential equations:

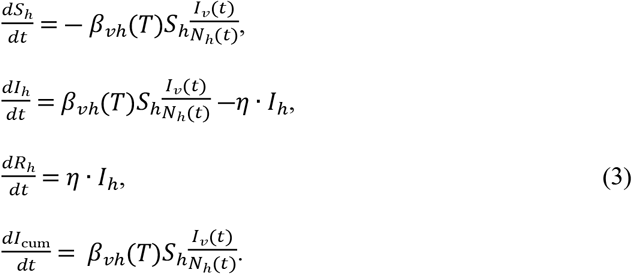

Here, *η* is the monthly recovery rate, calculated from the average infectious period in humans. Assuming an average infectious period of approximately 5 days [15], the recovery rate is *η* = 1/(5/30) = 6 per month.

### Dengue-Informed Neural Network (DINN)

Physics-Informed Neural Networks (PINNs) are data-driven frameworks designed to solve forward and inverse problems governed by differential equations [39]. Recent studies have demonstrated that PINN-based methods can accurately fit epidemiological data and infer time-varying parameters in compartmental-disease models [28-35]. Building on these advances, we propose a Dengue-Informed Neural Network (DINN) that jointly fits epidemiological observations and infers the time-varying level of infectious mosquitoes, *I*_*v*_ (*t*).

The DINN comprises two independent deep neural networks, 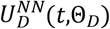 and 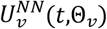,both of which take time *t* (in months) as the sole input:

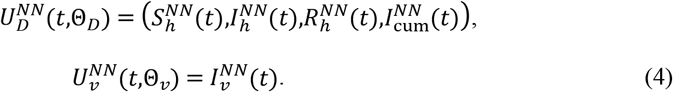

where 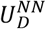 approximates the solution of the reduced SI-SIR system, while 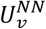 estimates the unobserved level of infectious mosquitoes. Both networks are implemented as fully connected feed-forward architectures with five hidden layers of 64 neurons each, using the hyperbolic tangent activation function [42].

The two networks in the DINN are trained jointly through a composite loss function that enforces agreement with both the epidemiological observations and the governing dynamical equations (Fig 1):

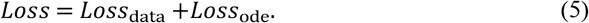

Minimizing this loss yields the optimal parameters Θ_*D*_ and Θ_*v*_, enabling simultaneous data fitting and inference of the time-varying infectious mosquito dynamics (*I*_*v*_(*t*)).

To ensure accuracy for fitting cumulative number of reported cases and derived number of new cases, the data-fitting loss function is defined as:

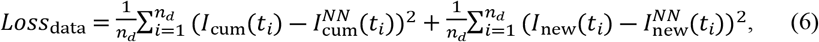

where (*n*_*d*_ = 136 is the number of monthly observations. To ensure consistency with transmission dynamics, deviation from the governing differential equations is penalized through an ODE residual loss function:

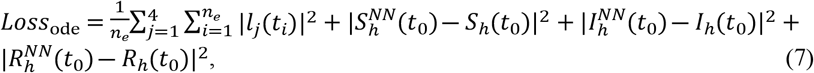

where *n*_*e*_ (= *n*_*d*_) denotes the number of residual points. The residuals *l*_*j*_(*t*_*i*_) correspond to those of the four differential equations:

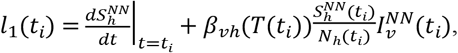

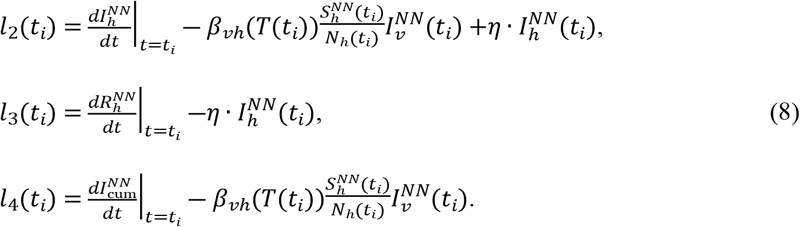

Initial conditions are set as *I*_*h*_(*t*_0_) = *I*_cum_(*t*_0_), *S*_*h*_(*t*_0_) = *N*_*h*_(*t*_0_) ― *I*_*h*_(*t*_0_), and *R*_*h*_(*t*_0_) = 0. Training was first performed using the Adam optimizer for 40,000 iterations, followed by refinement with the L-BFGS optimizer for up to 50,000 iterations. All computations were implemented in Python using the PyTorch deep learning framework [41]. Through inverse modelling with DINN, we obtained a continuous reconstruction of the latent mosquito infection dynamics *I*_*v*_(*t*) from reported monthly new (*I*_new_(*t*_*i*_)) and cumulative (*I*_cum_(*t*_*i*_)) cases.

To evaluate inference accuracy and biological consistency, the DINN-inferred mosquito infection time series *I*_*v*_(*t*) was substituted into the reduced SI-SIR model while keeping all other parameters fixed. The resulting ODE system was then solved numerically using a standard Python ODE solver. Agreement between the numerical solutions and the DINN-predicted epidemic trajectories was used to validate the inferred mosquito dynamics. Simulated monthly new *I*_new_(*t*), and cumulative *I*_cum_(*t*) cases were compared with observed data for all 15 countries using mean squared error (MSE) and coefficients of determination (*R*^2^).

### Recurrent Neural Networks for Mosquito Infection Forecasting

Using reconstructed mosquito infection dynamics from DINN, evaluated at monthly time points *I*_*v*_(*t*_*i*_), we developed a recurrent neural network to forecast mosquito infection levels. Specifically, lagged sequences of DINN-inferred mosquito infection *I*_*v*_(*t*_*i*_), together with lagged temperature and precipitation, were used to predict mosquito infection levels in the subsequent month.

Previous studies have provided strong empirical evidence for multi-month lagged effects of climate on arboviral transmission. Lowe et al. [55] demonstrated that the impact of drought conditions on dengue outbreaks can persist for up to five months. Similarly, Caldwell et al. [14] reported that the optimal predictive lags for dengue and chikungunya incidence consistently clustered at three to four months across ecologically and culturally diverse sites in Ecuador and Kenya, with statistically significant lag effects extending to five months in some locations. To capture this empirically observed range of delayed climate responses across heterogeneous settings, we adopted a unified five-month lag window.

Accordingly, the forecasting models used lagged DINN-inferred mosquito infection levels *I*_*v*_(*t*_*i*―1_),…, *I*_*v*_(*t*_*i*―5_), together with five-month lagged temperature *T*(*t*_*i*―1_),…, *T*(*t*_*i*―5_) and precipitation *P*(*t*_*i*―1_), …, *P*(*t*_*i*―5_), as input variables [14]. The target variable was the mosquito infection level in the subsequent month, *I*_*v*_(*t*_*i*_).

We considered three recurrent architectures: a vanilla (Elman-type) recurrent neural network (RNN), a gated recurrent unit (GRU), and a long short-term memory (LSTM) network [39]. Each model comprised two hidden layers with 32 neurons per layer. DINN-inferred mosquito infection data from January 2014 to April 2023 were used for training, while data from May 2023 to April 2025 were reserved for validation. To identify the optimal model for each country, we performed a grid search over learning rates {0.01, 0.005, 0.001}, training epochs {2000, 3000, 4000, 5000}, and batch sizes {16, 32, 64}. Model performance was evaluated using test-set mean squared error (MSE) and the coefficient of determination (*R*^2^), and the best-performing model was selected for each country (Supplementary Materials S1 Table).

Twenty-four-month forecasts of infected mosquitoes generated by the country-specific optimal recurrent neural networks, together with projected temperature and population data, were then integrated into the reduced SI-SIR model. Subsequent twenty-four-month forecasts of monthly dengue incidence was obtained by numerically solving the system (reduced SI-SIR model) using the odeint function from the SciPy library.

Finally, to quantify the contribution of individual predictors to model performance, we applied Shapley Additive Explanations (SHAP) analysis to the ensemble of country-specific optimal forecasting models. SHAP summary plots illustrate the distribution of feature contributions, with the horizontal axis representing SHAP values. In addition, to quantitatively assess the influence of climatic variables, we calculated the mutual information (MI) between time-lagged temperature or precipitation and infected mosquito abundance *I*_*v*_ (*t*) (Supplementary Materials, S1A Text). To assess whether the estimated MI reflects statistically significant dependence rather than random association, we further employed a permutation-based significance test under a null model (Supplementary Materials, S1B Text).

## Results

### Data Fitting and Parameter Inference

We trained the DINN using the reduced SI-SIR transmission model together with monthly dengue surveillance data *I*_new_(*t*_*i*_), human population size *N*_*h*_(*t*_*i*_), and monthly mean temperature *T*(*t*_*i*_) from 15 countries. Fig 2 shows the fitted monthly dengue incidence *I*_new_ (*t*) alongside the DINN-inferred time series of infected mosquitoes *I*_*v*_(*t*). The corresponding fits to cumulative monthly cases *I*_cum_(*t*) are shown in S2 Fig.

**Fig 2.**
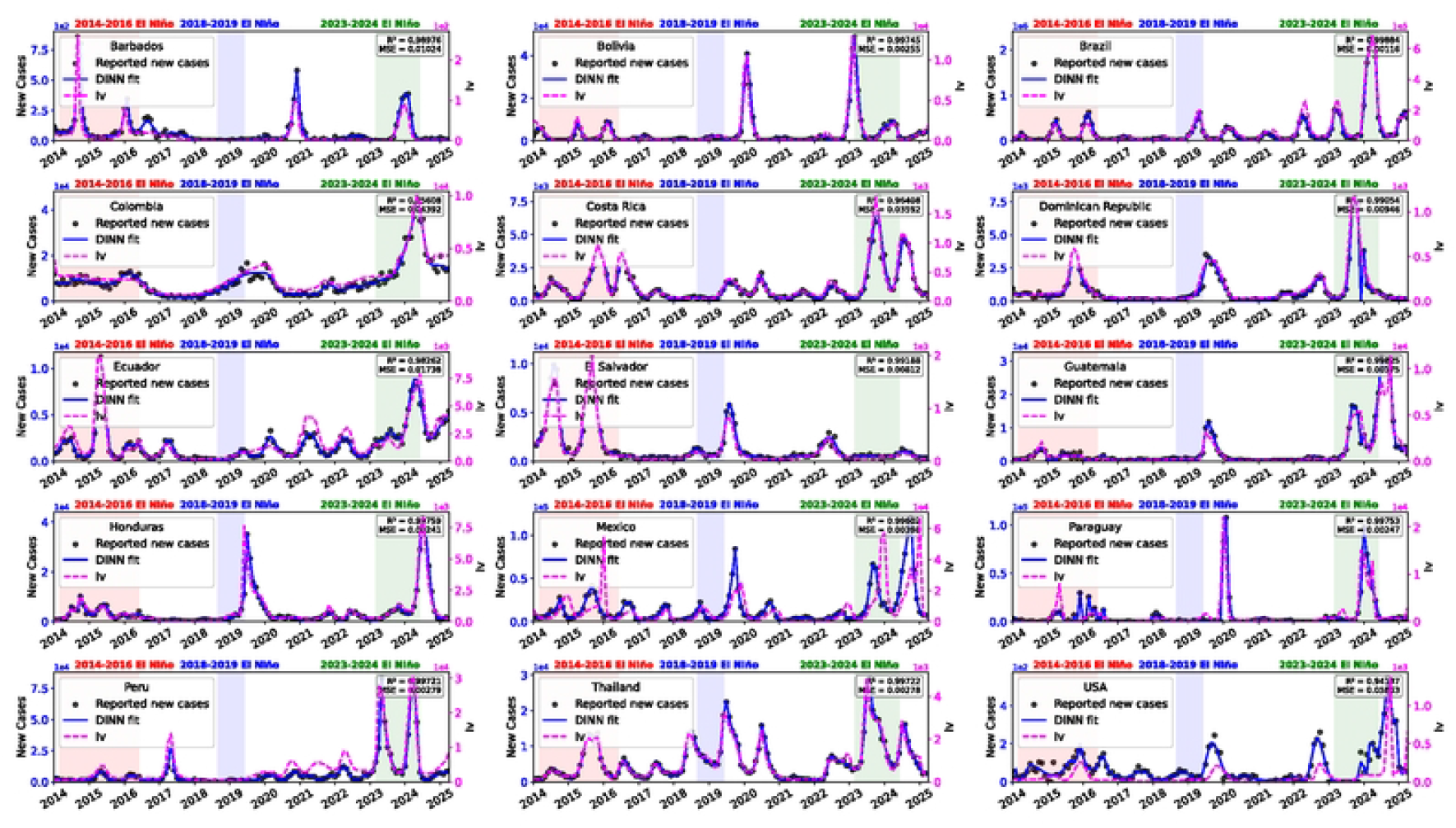
Data fitting and parameter inference results for dengue outbreaks in 15 countries using the DINN framework. Black dots represent observed monthly newly reported cases, and blue curves indicate the corresponding DINN-fitted trajectories. Pink curves denote the inferred time-varying abundance of infected mosquitoes (*I*_*v*_). Shaded regions indicate periods associated with El Niño events.

Across all countries, DINN successfully captured the multi-wave outbreak patterns characteristic of dengue epidemics and achieved excellent agreement with observed data (Fig 2), with coefficients of determination exceeding *R*^2^ > 0.94 and mean squared error *MSE* < 0.06 for monthly reported cases. Shaded regions indicate periods associated with El Niño events during 2014-2016, 2018-2019, and 2023-2024. During these intervals, the inferred mosquito infection dynamics exhibited one or multiple outbreak waves, with timing closely aligned with El Niño occurrence, suggesting that large-scale climatic variability plays an important role in driving dengue epidemic fluctuations. Notably, although only reported human incidence and temperature data were used for model training, DINN reconstructed biologically plausible mosquito infection dynamics *I*_*v*_(*t*), demonstrating its capacity to infer unobserved vector processes from limited epidemiological observations.

### Model Verification

To assess the biological consistency of the DINN-inferred mosquito infection dynamics, we incorporated the estimated *I*_*v*_(*t*) as a time-varying input into the reduced SI-SIR model. We then numerically simulated monthly new dengue cases *I*_new_(*t*) (Fig 3) and cumulative cases *I*_cum_(*t*) (S3 Fig), and compared with observed data across all 15 countries. With the exception of the United States (*R*^2^ = 0.466, MSE = 0.534), the numerical solutions closely reproduced the observed epidemic trajectories for all other countries (*R*^2^ > 0.88 and MSE < 0.12; Fig 3). This strong agreement confirms that the DINN-derived *I*_*v*_(*t*) trajectories are compatible with the underlying transmission dynamics. Overall, these results demonstrate that the inferred vector infection dynamics provide a coherent mechanistic explanation for the observed dengue epidemics.

**Fig 3.**
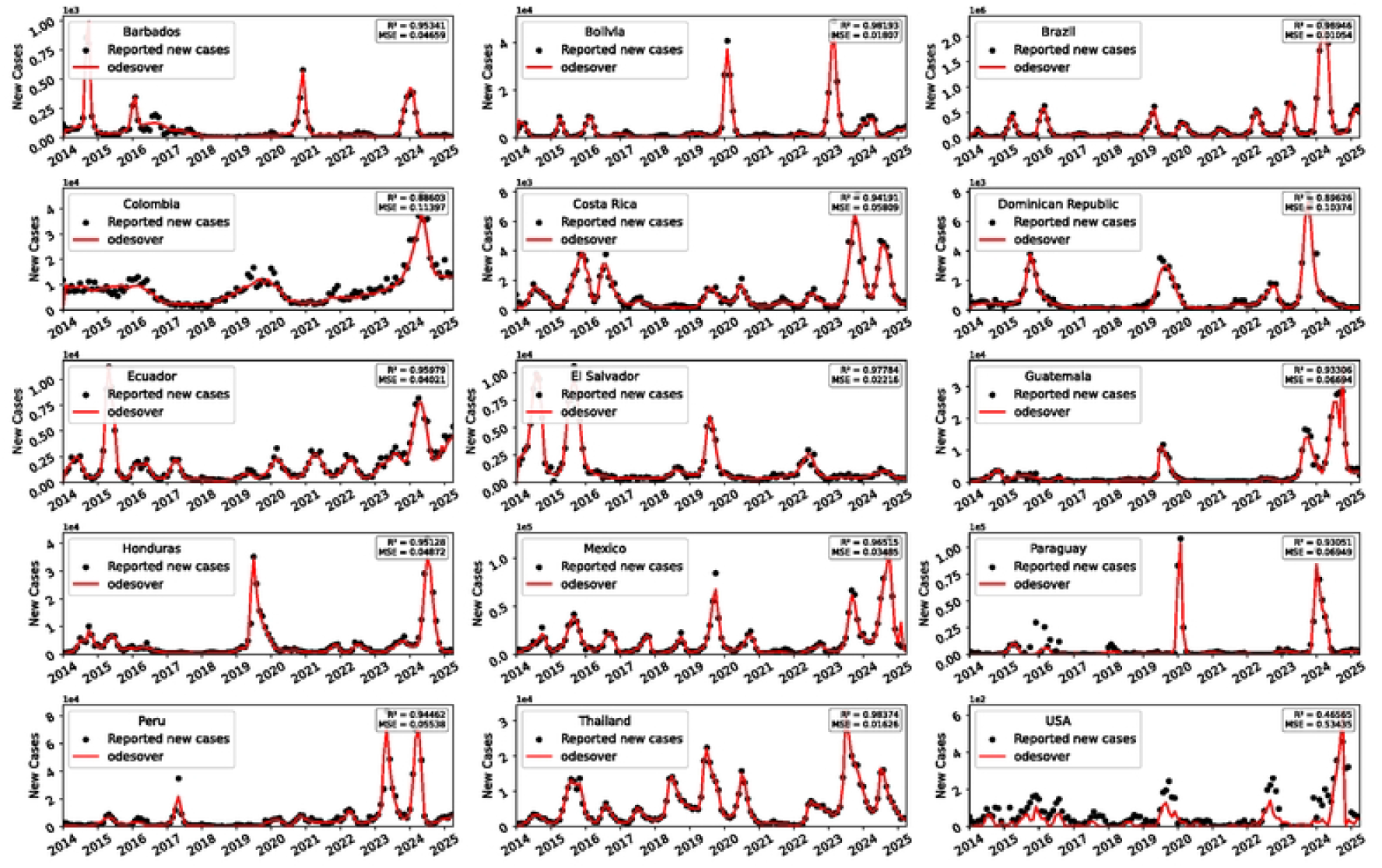
Epidemic trajectories generated by substituting the DINN-inferred infected mosquito dynamics (*I*_*v*_) into the reduced SI-SIR model. Black dots represent observed monthly newly reported cases, and the red curves show the corresponding numerical solutions of model (3).

### Forecasting Mosquito and Dengue Dynamics

Using the best-performing recurrent model for each country, we generated twenty-four-month forecasts of infected mosquitoes *I*_*v*_(*t*) from May 2023 to April 2025 (red curves in Fig 4). The MSE (mean square error) Loss of each optimal recurrent model decreased toward zero as training progressed (i.e., with increasing of epochs; S4 Fig), indicating that the average squared difference between DINN-inferred mosquito infection levels and recurrent model predictions was small.

**Fig 4.**
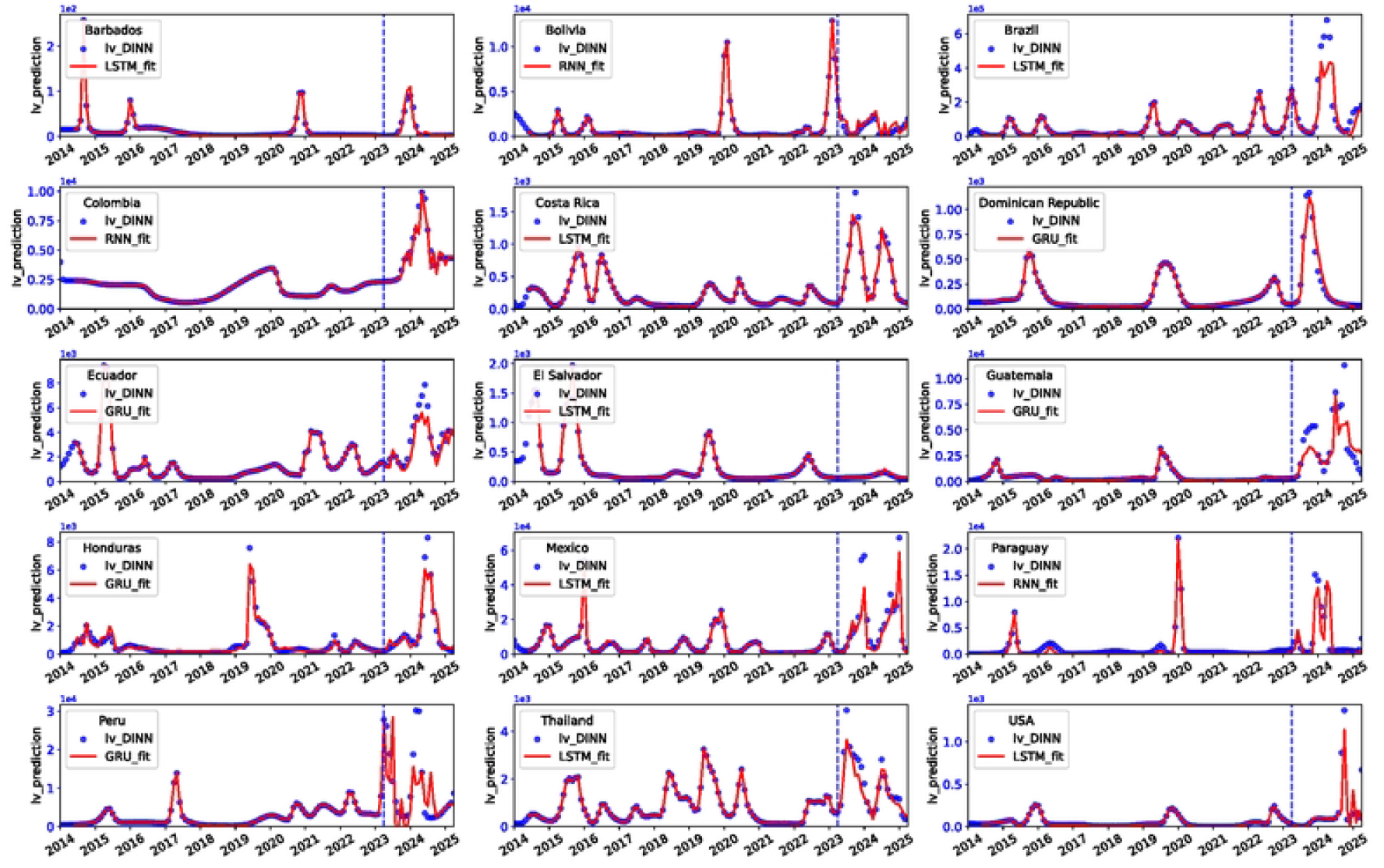
Twenty-four-month forecasts (May 2023-April 2025) of infected mosquitoes (*I*_*v*_) for each country generated using the best-performing model (RNN, GRU, or LSTM). Blue dots denote DINN-inferred mosquito infection data used for model training and testing. Red curves indicate fitted values during the training period (left of the vertical line) and forecasts during the prediction period (right of the vertical line).

Forecast performance closely reflected the test *R*^2^ and MSE values reported in Table 1. Countries with high test *R*^2^ and low test MSE (e.g., Barbados, Costa Rica, Thailand, the Dominican Republic, and Honduras) exhibited accurate and well-calibrated mosquito infection forecasts, capturing both seasonal peaks and interannual variation, including El Niño–associated surges during late 2023 to early 2024. In contrast, countries with low test *R*^2^ values (e.g., Peru, Bolivia, and Guatemala) showed poorer forecast alignment, likely reflecting data limitations or unmodelled local drivers of transmission (Fig 4; Table 1).

**Table 1.** Optimal hyperparameter settings and predictive performance of the selected forecasting models during training and testing for each country.

| Country | Model | Learning rate | Number of epochs | Batch size | Train MSE | Test MSE | Train R<sup>2</sup> | Test R<sup>2</sup> |
| --- | --- | --- | --- | --- | --- | --- | --- | --- |
| Barbados | LSTM | 0.005 | 3000 | 16 | 4.78E-07 | 0.001289 | 0.999963 | 0.882615 |
| Bolivia | RNN | 0.001 | 3000 | 32 | 1.06E-05 | 0.002417 | 0.999599 | 0.132022 |
| Brazil | LSTM | 0.001 | 3000 | 64 | 2.99E-05 | 0.026447 | 0.9955 | 0.692957 |
| Colombia | RNN | 0.001 | 2000 | 32 | 1.36E-05 | 0.006551 | 0.997818 | 0.869953 |
| Costa Rica | LSTM | 0.001 | 3000 | 32 | 6.48E-06 | 0.005563 | 0.999472 | 0.926842 |
| Dominican Republic | GRU | 0.005 | 2000 | 64 | 2.66E-05 | 0.005413 | 0.997977 | 0.94229 |
| Ecuador | GRU | 0.005 | 2000 | 16 | 1.37E-06 | 0.011727 | 0.999954 | 0.712649 |
| El Salvador | LSTM | 0.005 | 5000 | 32 | 3.32E-07 | 0.000107 | 0.999992 | 0.599964 |
| Guatemala | GRU | 0.001 | 3000 | 16 | 8.77E-06 | 0.032732 | 0.996129 | 0.478717 |
| Honduras | GRU | 0.01 | 3000 | 16 | 0.000875 | 0.007918 | 0.938781 | 0.881843 |
| Mexico | LSTM | 0.001 | 3000 | 64 | 3.88E-05 | 0.019098 | 0.996287 | 0.737783 |
| Paraguay | RNN | 0.005 | 4000 | 16 | 0.000489 | 0.017897 | 0.963746 | 0.577471 |
| Peru | GRU | 0.01 | 2000 | 16 | 1.77E-06 | 0.069548 | 0.999864 | 0.198932 |
| Thailand | LSTM | 0.005 | 2000 | 32 | 5.45E-07 | 0.009831 | 0.999974 | 0.833479 |
| USA | LSTM | 0.005 | 3000 | 64 | 2.51E-07 | 0.021642 | 0.999869 | 0.592122 |

The predicted mosquito infection time series were subsequently used as inputs to the reduced SI-SIR model to simulate future dengue incidence. Forecasts of monthly newly reported and cumulative cases for all 15 countries are shown as red lines in Fig 5 and S5 Fig, respectively. With the exception of the United States, countries with strong mosquito-forecast performance (high test *R*^2^ and low test MSE) also exhibited close agreement between predicted and observed dengue incidence, for both newly reported and cumulative cases. These results highlight the reliability of the integrated forecasting framework when the underlying mosquito infection dynamics are accurately predicted.

**Fig 5.**
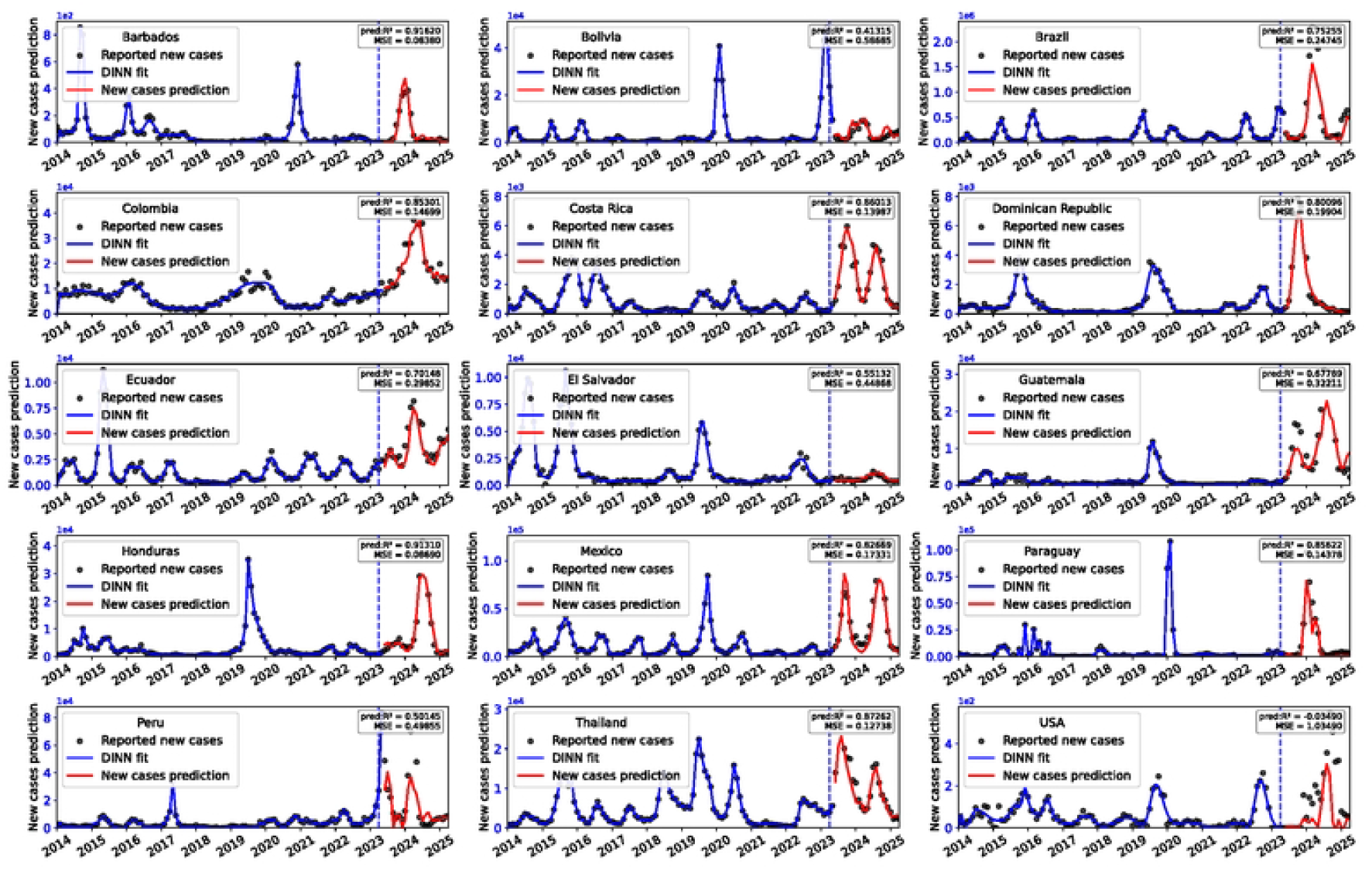
Twenty-four-month forecasts (May 2023-April 2025) of monthly newly reported dengue cases using predicted *I*_*v*_ from each country’s best-performing model. Black dots represent observed cases. Blue curves to the left of the vertical line indicate DINN-fitted result, whereas red curves to the right show forecasts generated by model (3).

### Interpretability Analysis

Fig 6 presents the mean absolute SHAP values for all predictor variables. Across countries, lagged mosquito infection variables (*I*_*v*_lag1_, *I*_*v*_lag2_, *I*_*v*_ lag3_) consistently exhibited the highest importance scores, indicating that short-term historical dynamics of infected mosquitoes are the dominant drivers of model predictions. In contrast, temperature and precipitation contributed relatively less, suggesting that although climatic conditions modulate mosquito abundance, their effects are secondary to endogenous vector population dynamics.

**Fig 6.**
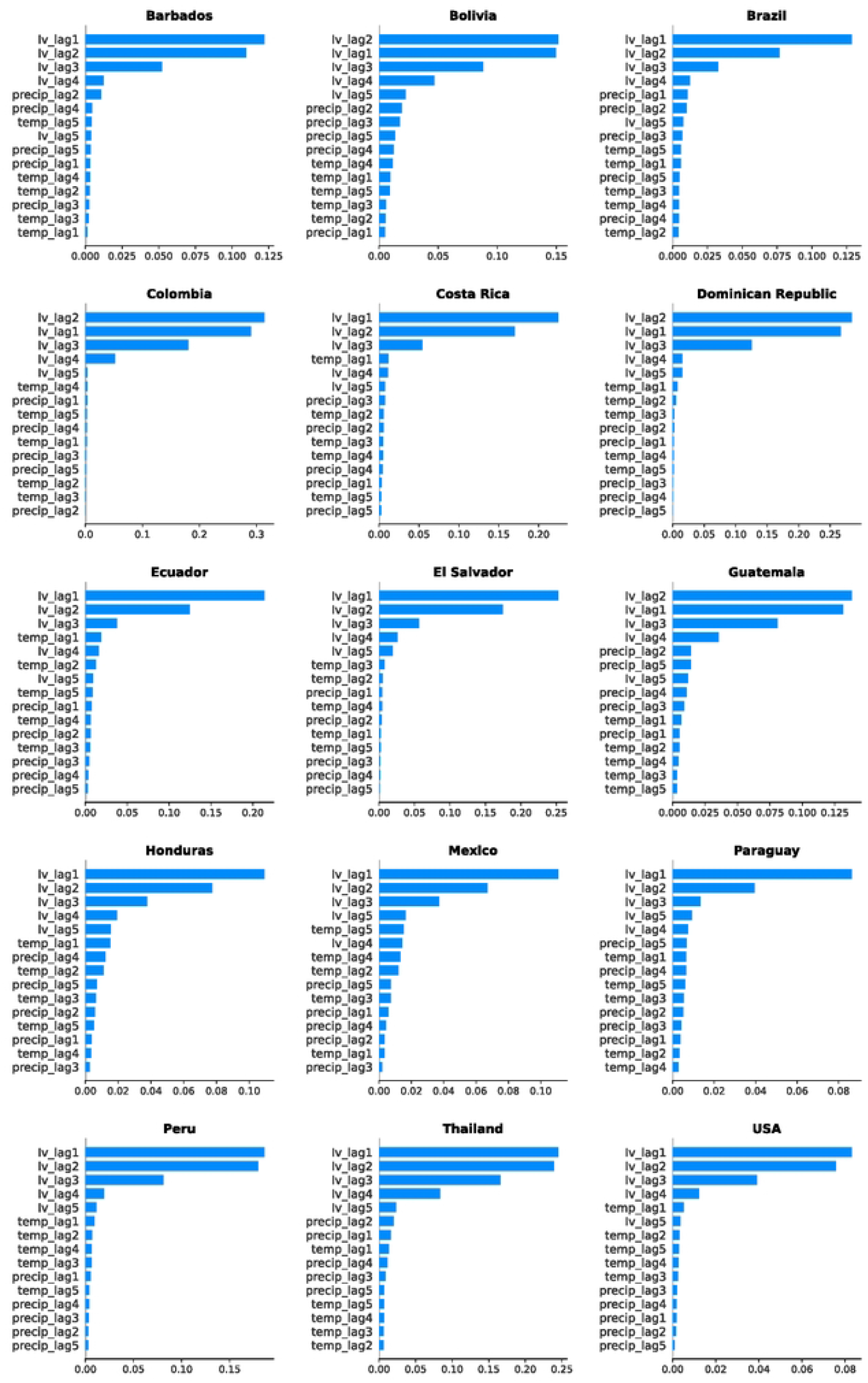
SHAP feature-importance plots for the optimal prediction models across all 15 countries.

SHAP summary plots (Fig 7) further illustrate the distributions of feature contributions, with the horizontal axis representing SHAP values. *I*_*v*_lag1_ exhibited a strong positive association with *I*_*v*_(*t*), with higher feature values (red points) yielding positive SHAP contributions. In contrast, *I*_*v*_lag2_ showed a negative association, with larger values corresponding to negative SHAP contributions, reflected by red points clustering in the negative SHAP region and blue points in the positive region (Fig 7; S6, S7 Figs). As shown in Fig 7 and S8 Fig, *I*_*v*_lag3_ was positively associated with *I*_*v*_(*t*), with red points concentrated in the positive SHAP region and blue points in the negative region.

**Fig 7.**
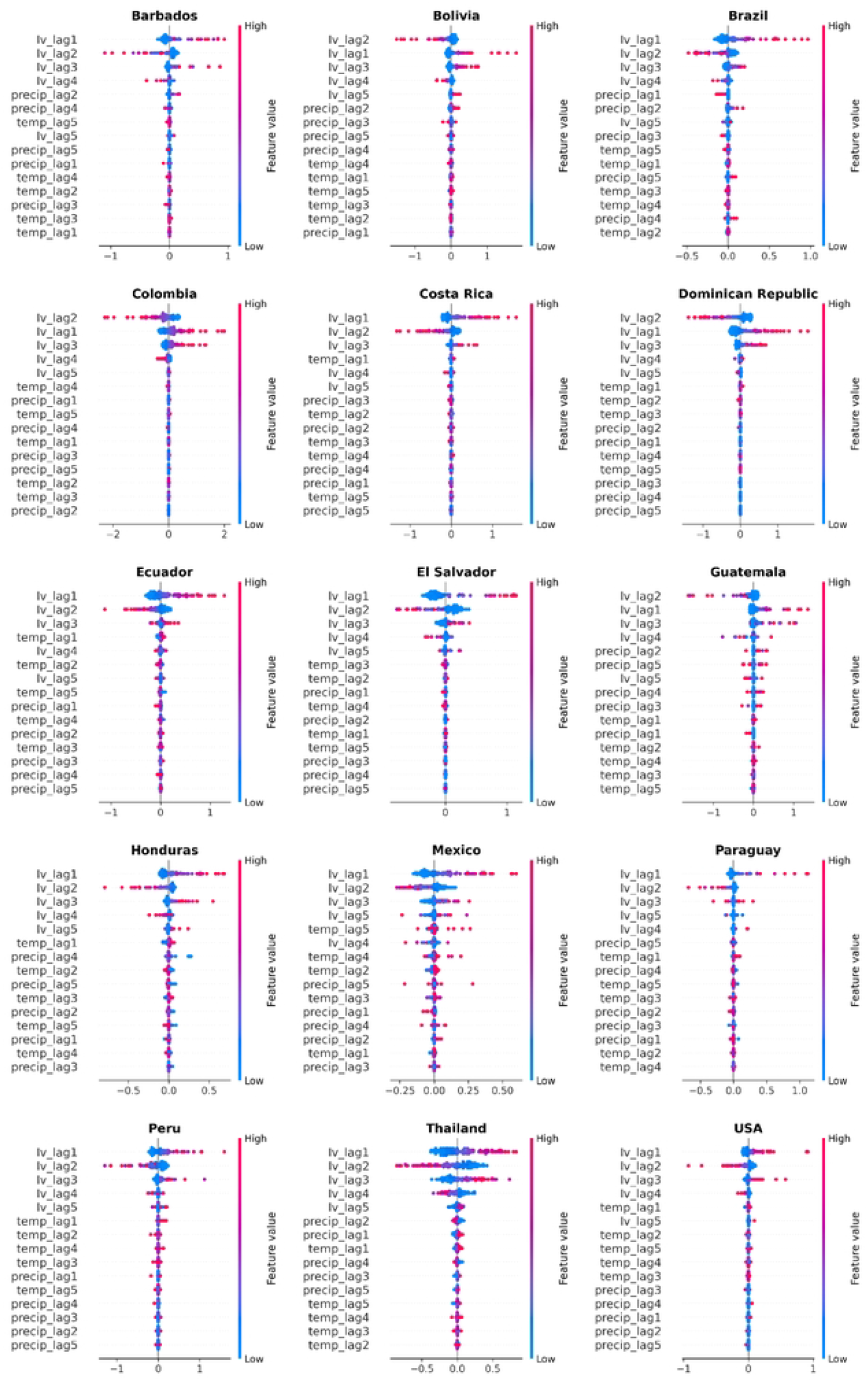
SHAP summary plots showing the distributions of feature contributions for the optimal prediction models across all 15 countries.

Substantial inter-country heterogeneity was also observed. For example, *I*_*v*_lag1_ exerted a strong predictive influence in Barbados, Brazil, and Costa Rica, whereas its importance was considerably lower in Bolivia, the Dominican Republic, and Guatemala (Fig 6). This heterogeneity highlights geographic variation in the mechanisms governing mosquito infection dynamics.

Mutual information analysis further quantified the influence of climatic variables on mosquito infection. Lagged temperature was correlated with infected mosquito abundance in all countries and exhibited statistically significant associations in Bolivia, Colombia, Costa Rica, EI Salvador, Guatemala, and Peru (p-value < 0.05; S9a, b Fig). Except for Costa Rica, lagged precipitation exhibited a dependency relationship with infected mosquitoes across all countries (S9c Fig). In particular, lagged precipitation showed statistically significant associations in Brazil, Colombia, Guatemala, Peru, and the Dominican Republic (p-value<0.05; S9d Fig).

## Discussion

Vector control remains central to dengue prevention, yet predicting outbreaks of infected mosquitoes is challenging because real-time entomological surveillance data are rarely available. To address this gap, we developed a Dengue-Informed Neural Network (DINN) that integrates a reduced SI-SIR transmission model with physics-informed neural networks to infer unobserved mosquito infection dynamics from reported dengue case data. Applying DINN to monthly dengue surveillance data from 15 countries, we fitted observed dengue incidence and reconstructed latent time series of infected mosquitoes.

The DINN framework establishes a bidirectional coupling between deep learning and the reduced SI-SIR transmission model. Neural networks infer latent mosquito infection dynamics directly from human case data, while the governing differential equations impose mechanistic constraints that regularise learning (Fig 1). Reverse simulation using an ODE solver confirmed that the inferred mosquito trajectories are biologically consistent and capable of reproducing observed epidemic dynamics. Across countries, DINN captured characteristic multi-wave dengue outbreaks and revealed substantial regional heterogeneity in inferred mosquito infection dynamics (Fig 2).

Building on these reconstructed vector dynamics, we developed recurrent neural network– based forecasting models to predict mosquito infection levels up to 24 months ahead and subsequently projected future dengue incidence. Overall, the integrated framework demonstrated strong predictive performance in most countries and provided an interpretable, mechanistically grounded basis for dengue early warning and decision support.

### The Role of the Reduced SI-SIR Model: Limitations of Direct Incidence Forecasting

A key methodological contribution of this study is the use of a reduced SI-SIR model in which the number of infected mosquitoes, *I*_*v*_(*t*), is treated as a latent, time-varying input rather than as an explicitly modelled compartment. This design is essential for vector-borne diseases such as dengue [15,17], where mosquito population sizes and infection status are rarely observed and are strongly influenced by environmental variability.

Existing PINN-based hybrid modelling studies have largely focused on directly transmitted diseases, such as COVID-19, whereas applications to vector-borne diseases remain limited [53]. Compared with traditional mechanistic models, PINNs provide greater flexibility for learning unknown or time-varying transmission processes directly from data. At the same time, relative to purely data-driven deep learning approaches, PINNs retain mechanistic interpretability through the incorporation of governing equations explicitly as physical constraints [30]. Many previous hybrid frameworks further rely on ODE solvers to extrapolate epidemic trajectories using fixed parameters estimated at the end of the training period [32,35]. Although such assumptions may be reasonable for diseases with relatively stable transmission rates, they are poorly suited to vector-borne diseases, which are characterised by pronounced environmental seasonality and recurrent epidemic waves.

By treating *I*_*v*_(*t*) as key latent variable, the reduced SI-SIR model preserves essential vector-host coupling while avoiding strong and often unverifiable assumptions about mosquito demography and population dynamics. Inferring and forecasting *I*_*v*_(*t*) therefore resolves a central limitation of classical dengue models and provides a more robust, flexible,and interpretable basis for predicting human incidence.

### The Role of Climate: Temperature, Rainfall, and Large-Scale Climate Variability

Climatic variables influence dengue transmission indirectly by shaping mosquito life-history traits, development rates, survival, and population density [47-49]. Because direct observations of mosquito populations are scarce, most existing early warning systems rely primarily on climatic time series to predict dengue incidence [50-52]. In our framework, climate effects are represented both implicitly in the DINN-inferred mosquito infection trajectories and explicitly through their inclusion as predictors in mosquito forecasting models.

Our results indicate that short-term lagged mosquito infection history consistently dominates predictive performance (Fig 6), reflecting strong temporal autocorrelation and endogenous vector population dynamics. Nevertheless, temperature and precipitation exhibited statistically significant associations with mosquito infection levels in many countries, as confirmed by mutual information analysis. These findings support the view that climate primarily acts as a modulating, rather than a direct driving, force in shaping short-term dengue dynamics.

Importantly, the inferred mosquito infection trajectories displayed pronounced increases during El Niño periods (2014-2016, 2018-2019, and 2023-2024), with outbreak timing closely aligned across multiple regions. This synchrony highlights El Niño as a major large-scale climatic driver capable of coordinating dengue risk across geographically distant settings, consistent with previous studies [44–46]. Incorporating both local climate variables and large-scale climate oscillations is therefore essential for developing robust and transferable dengue early warning systems.

### Geographic Heterogeneity in Model Performance and Predictive Reliability

Substantial geographic heterogeneity was observed in both inference and forecasting performance across countries. Countries such as Barbados, Costa Rica, Thailand, and the Dominican Republic showed high predictive accuracy, with forecasts successfully capturing seasonal cycles, interannual variability, and El Niño-associated surges (Fig 2; Fig 5). These countries are characterised by relatively regular transmission patterns and more consistent surveillance, facilitating reliable inference of underlying vector dynamics.

In contrast, predictive performance was lower in countries such as Bolivia, Peru, and the United States. This likely reflects a combination of under-reporting, spatial heterogeneity masked by national-level aggregation, and unmodelled local drivers such as vector control interventions, healthcare access, and population mobility. These factors introduce additional sources of variability that are not captured by the current modelling framework. Overall, these findings highlight that while the DINN framework is broadly transferable across diverse epidemiological contexts, its performance depends critically on data quality and on the extent to which national-level surveillance time series accurately reflect local transmission processes. Future extensions incorporating subnational data and additional covariates may further enhance predictive reliability in heterogeneous settings.

### Limitations and Concluding Remarks

Despite its advantages, several limitations should be acknowledged. First, dengue incidence and climate data were aggregated at national and monthly scales, resulting in coarse spatiotemporal resolution that may obscure important local heterogeneity in transmission dynamics. Second, ecologically relevant factors such as humidity, vegetation indices, and land-use patterns were not included due to data limitations [54]. Third, the reduced SI-SIR model does not explicitly account for socio-behavioural factors, public health interventions, or vaccination coverage, all of which can substantially modify transmission dynamics.

Future work should seek to integrate higher-resolution epidemiological and environmental data, incorporate additional ecological and social drivers, and explicitly model intervention strategies. Extending the proposed framework to other vector-borne diseases would further assess its generalisability and practical utility. Despite these limitations, the DINN framework provides a flexible, interpretable, and mechanistically grounded approach for inferring unobserved vector dynamics and linking environmental variability to human disease outbreaks. By explicitly modelling infected mosquitoes as a latent yet forecastable state, this framework addresses key shortcomings of both traditional mechanistic models and purely data-driven approaches, and offers a promising foundation for the development of robust early warning systems for dengue and other mosquito-borne diseases.

## Acknowledgements

M.S. is supported by the National Natural Science Foundation of China (No. 32371555), Natural Science Foundation of Anhui Province (No. 2308085MA09) and the Fundamental Research Funds for the Central Universities (No. JZ2024HGTG0278). C.H. acknowledges support from the South African Research Chair Initiative (National Research Foundation of South Africa, grant 89967).

## Data Availability

Monthly dengue case data were obtained from the World Health Organization (WHO) [1]. Climatic variables, including temperature and precipitation, were retrieved from the National Oceanic and Atmospheric Administration (NOAA) [36], while population data were obtained from the World Bank and Worldometer [37,38]. All processed datasets and model-generated outputs supporting the figures and tables in this study (including DINN-inferred monthly mosquito infection dynamics for all 15 countries and forecasting results), are publicly available at: https://doi.org/10.5281/zenodo.18521916. The complete implementation of the Dengue-Informed Neural Network (DINN), including its two component neural networks, the reduced SI–SIR model, and all candidate recurrent forecasting models (RNN, GRU, and LSTM), as well as scripts for data preprocessing, model training, validation, and simulation, is available at: https://github.com/Caojiayue/DINN. All trained models, including the optimised DINN networks, inferred reduced SI–SIR model parameters, and country-specific best-performing forecasting models, are archived at: https://doi.org/10.5281/zenodo.18521916. These resources enable full reproducibility of the analyses and results reported in this study.

## Author contributions

**Conceptualization:** Jiayue Cao, Ziqiang Cheng, Min Su, Cang Hui

**Methodology:** Jiayue Cao, Ziqiang Cheng, Min Su, Cang Hui

**Supervision:** Min Su, Cang Hui

**Validation:** Jiayue Cao

**Visualization:** Jiayue Cao, Min Su

**Writing-original draft:** Jiayue Cao, Min Su

**Writing-review & editing:** Min Su, Cang Hui

## Supplementary Online Materials

**S1 Fig**. Monthly temperature and precipitation time series for all 15 countries.

**S2 Fig**. DINN fitting results for cumulative reported dengue cases across 15 countries. Black dots represent observed cumulative case data, and blue curves denote the DINN-based best-fit trajectories.

**S3 Fig**. Numerical solutions of the reduced SI-SIR model obtained by incorporating the DINN-inferred infected mosquito time series *I*_*v*_(*t*). Black dots represent observed cumulative reported cases, and red curves show the corresponding solutions of model (3).

**S4 Fig**. MSE (mean squared error) loss of the optimal prediction model for each country during training.

**S5 Fig**. Twenty-four-month forecasts (May 2023-April 2025) of cumulative reported dengue cases for each country generated using the best-performing prediction model. Black dots denote observed cumulative cases. Blue curves to the left of the vertical line indicate DINN-fitted results, whereas red curves to the right represent forecasts produced by model (3). The coefficients of determination *R*^2^ and mean squared errors (MSE) summarise forecast performance.

**S6 Fig**. SHAP dependence plot for the most important predictor variable.

**S7 Fig**. SHAP dependence plot for the second most important predictor variable.

**S8 Fig**. SHAP dependence plot for the third most important predictor variable.

**S9 Fig**. Heatmap of mutual information (MI) values between predictors and infected mosquito abundance, with corresponding statistical significance.

**S1 Table**. Hyperparameter settings and predictive performance of all candidate prediction models (RNN, GRU, and LSTM) during training and testing.

**S1 Text**. Description of the mutual information estimation method and permutation-based significance testing procedure.

